# White matter microstructural changes in short-term learning of a continuous visuomotor sequence

**DOI:** 10.1101/2020.10.02.324004

**Authors:** Stéfanie A. Tremblay, Anna-Thekla Jäger, Julia Huck, Chiara Giacosa, Stephanie Beram, Uta Schneider, Sophia Grahl, Arno Villringer, Christine L. Tardif, Pierre-Louis Bazin, Christopher J. Steele, Claudine J. Gauthier

## Abstract

Efficient neural transmission is crucial for optimal brain function, yet the plastic potential of white matter (WM) has long been overlooked. Growing evidence now shows that modifications to axons and myelin occur not only as a result of long-term learning, but also after short training periods. Motor sequence learning (MSL), a common paradigm used to study neuroplasticity, occurs in overlapping learning stages and different neural circuits are involved in each stage. However, most studies investigating short-term WM plasticity have used a *pre-post* design, in which the temporal dynamics of changes across learning stages cannot be assessed. In this study, we used multiple magnetic resonance imaging (MRI) scans at 7 Tesla to investigate changes in WM in a group learning a complex visuomotor sequence (LRN) and in a control group (SMP) performing a simple sequence, for 5 consecutive days. Consistent with behavioral results, where most improvements occurred between the two first days, structural changes in WM were observed only in the early phase of learning (d1-d2), and in overall learning (d1-d5). In LRNs, WM microstructure was altered in the tracts underlying the primary motor and sensorimotor cortices. Moreover, our structural findings in WM were related to changes in functional connectivity, assessed with resting-state functional MRI data in the same cohort, through analyses in regions of interest (ROIs). Significant changes in WM microstructure were found in a ROI underlying the right supplementary motor area. Together, our findings provide evidence for highly dynamic WM plasticity in the sensorimotor network during short-term MSL.

## INTRODUCTION

The idea that structure determines function, and that function can modulate structure, is a well-known concept governing biology (Kohn et al., 2018). Just like any other organ in the body, the structure of the brain changes in response to changing demands in order to support new functions, in a process termed neuroplasticity (Zatorre et al., 2012). Synaptic changes have been the main focus of early plasticity studies (Rioult-Pedotti et al., 2000; Xu et al., 2009), yet recent research now indicates that plastic changes can also involve alterations to neurons, glial cells, and cerebral vessels (Sampaio-Baptista & Johansen-Berg, 2017; Tardif et al., 2016; Zatorre et al., 2012).

The plastic potential of white matter (WM), and the behavioral relevance of changes in the fiber tracts connecting neurons, has long been overlooked. However, growing evidence now shows modifications to astrocytes, microglia, and myelin-producing oligodendrocytes occur as a result of experience-dependent learning (Chorghay et al., 2018; Tardif et al., 2016). Changes to axons and myelin would lead to changes in conduction speed and thus more efficient information processing through optimized timing of neural transmission (Chorghay et al., 2018; Fields, 2015; Sampaio-Baptista & Johansen-Berg, 2017). Given the crucial role of efficient neural transmission for optimal brain function (Fields, 2015; Waxman, 1975), a deeper understanding of the ways in which WM can be altered by experience is of critical importance.

Motor sequence learning (MSL) tasks are a common paradigm used to study neuroplasticity (Doyon et al., 2009; O. Hikosaka et al., 2002; Nissen & Bullemer, 1987; Penhune & Steele, 2012) MSL occurs in overlapping stages that have been described by various models. As such, motor learning can be divided into an initial fast stage, where a large amount of improvement occurs in a short period of time, followed by a consolidation stage, which solidifies the gains in performance between training sessions, making them resistant to interference. In a final late/slow stage, the learned sequence is fine-tuned to optimize motor parameters such as force, timing and spatial accuracy (Dayan & Cohen, 2011; Doyon et al., 2002; Doyon & Benali, 2005; Avi Karni & Sagi, 1993; Korman et al., 2003; Luft & Buitrago, 2005). In each case, there is significant evidence from neuroimaging studies that different neural circuits are involved in each stage of learning (see Dayan & Cohen, 2011; Halsband & Lange, 2006 for reviews).

Studying neuroplasticity with magnetic resonance imaging (MRI) allows for the longitudinal investigation of functional and structural reorganization at the network (how are different brain regions connected to each other, i.e. whole-brain level) and microstructural levels (what properties do the fiber bundles that make up these connections have, i.e. voxel level), as whole-brain images can be obtained repeatedly (Tardif et al., 2016). Recent advances in MRI techniques and models, especially in the areas of connectivity and network theory, have created the opportunity for a better understanding of how brain architecture and network efficiency impact information processing and how these are modified through experience (Albert et al., 2009; Guye et al., 2010; Lewis et al., 2009). Changes in connectivity can be defined structurally, with diffusion-weighted imaging (DWI), a technique that probes WM microstructural organization through imaging the bulk motion of water molecules (Abdul-Kareem et al., 2011; Klein et al., 2019; Le Bihan et al., 2001), and functionally, through the measurement of spontaneous activity at rest and the temporal correlation of the blood-oxygen-level-dependent (BOLD) signal between brain regions (resting-state functional MRI) (Albert et al., 2009; Lewis et al., 2009).

In parallel, advances in hardware, such as the increasing use of ultra-high field MRI, and improved modeling approaches allow for the characterization and quantification of several properties of brain structures at finer spatial scales (Dumoulin et al., 2018; Frangou et al., 2020; Heidemann et al., 2012). These advances could allow us to bridge the gap between the knowledge gained from animal and human studies, and better define the mechanisms and time course at play when the brain is reshaped through experience (Sampaio-Baptista & Johansen-Berg, 2017; Tardif et al., 2016).

Although the greatest gains in performance occur during the initial stage of learning (Dayan & Cohen, 2011; Halsband & Lange, 2006; Savion-Lemieux & Penhune, 2005), early investigations of WM plasticity mainly focused on the effects of long-term training with cross-sectional studies in musicians and dancers with several years of training (Abdul-Kareem et al., 2011; Bengtsson et al., 2005; Giacosa et al., 2016, 2019; Hänggi;nggi et al., 2010). The cross-sectional nature of many of these studies does not allow characterization of the temporal dynamics of learning nor distinguishing training-induced changes from pre-existing differences in WM (e.g. Abdul-Kareem et al., 2011; Bengtsson et al., 2005; Giacosa et al., 2016, 2019; Hängginggi et al., 2010; Steele et al., 2012). More recently, longitudinal studies using DWI have shown that WM changes can occur at shorter timescales. For instance, changes to WM microstructure underlying the intraparietal sulcus were observed after 6 weeks of juggling training (Scholz et al., 2009) and another study reported such changes in the fornix as quickly as after a few hours of spatial learning in a car racing game (Hofstetter et al., 2013). However, most studies investigating short-term WM plasticity have used a *pre-post* design, or a single measurement at the end of learning, in which the temporal dynamics of changes across learning stages cannot be assessed (Hofstetter et al., 2013; Scholz et al., 2009; Steele et al., 2012). Moreover, the control group in some of these studies does not allow to distinguish changes due to sequence-specific learning from those due to motor execution (e.g. Scholz et al., 2009). In this study, we used multiple high-resolution MRI scans at 7T to investigate changes in WM across learning stages in a group learning a complex visuomotor sequence, and in a control group performing a simple sequence. Moreover, we investigated WM plasticity in regions of interest (ROIs) near areas of change in functional connectivity, assessed with resting-state fMRI (rs-fMRI) data in the same cohort (unpublished).

## METHODS

### Participants

Forty neurologically normal individuals of 21 to 30 years of age (M ± SD: 24.5 ± 2.44; 21 females) and without motor or correctible visual impairments were recruited from the participant database of the Max Planck Institute for Human Cognitive and Brain Sciences in Leipzig, Germany. All participants were task naïve prior to this study and right-handed according to the Edinburgh Handedness Inventory (M ± SD: 83.7 ± 16.9), except for one who was ambidextrous (EHI= 40). The majority of participants had no exceptional musical experience (M ± SD: 8.55 ± 8.83 years), but one participant self-identified as a professional musician and two as having advanced musical experience. Participants had an average sport experience of 5.83 ± 7.15 years and two participants self-identified as professional athletes. After ensuring the participants had no neurological conditions and no contraindication to MRI, they gave written informed consent according to the declaration of Helsinki. Participants were randomized into two groups: the experimental group (N=20), who learned a complex visuo-motor sequence, and the control group (N=20), who learned a simple repetitive visuo-motor sequence. One participant from the experimental group was excluded from this study due to a large signal drop in DWI data. Table 1 shows the demographic data for each group. After completion of the study, participants were financially compensated for their time. The study was approved by the ethics review board of the Medical Faculty of the University of Leipzig and all participants provided written informed consent according to the Declaration of Helsinki.

**Table 1.**
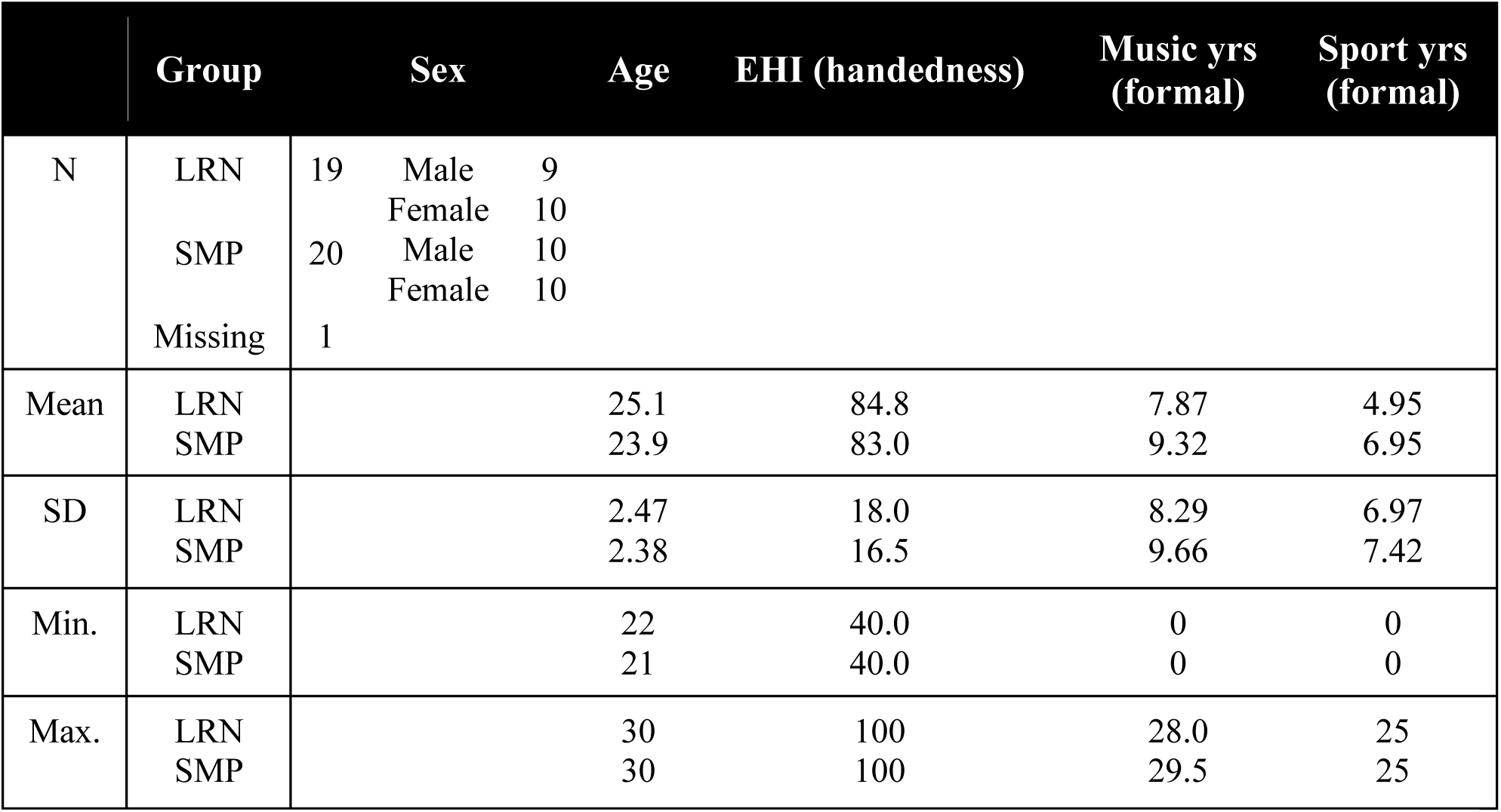
Demographic data in each group. LRN= experimental group; SMP= control group; SD= standard deviation; EHI= Edinburgh Handedness Inventory.

### Motor Sequence Learning Task

The sequential pinch force task (SPFT) is a complex visuomotor sequence learning task requiring fine force control (Camus et al., 2009; Krakauer et al., 2019), and was previously shown to result in short-term plastic changes in grey matter (Gryga et al., 2012). During the SPFT, participants hold a pressure sensor between the thumb and index finger of their right hand (Fig. 1a) and are required to exert force on the sensor in order to match the height of a moving reference bar (REF; blue on Fig. 1b) displayed on a computer screen. Another moving bar (FOR; yellow), representing the amount of pressure they are exerting on the device, is also displayed on the screen to provide visual feedback. The device samples force continuously throughout the task at a rate of 80 Hz. The change in height of the REF bar follows one of two specific sequences, as illustrated in Fig. 1 (c).

**Figure 1.**
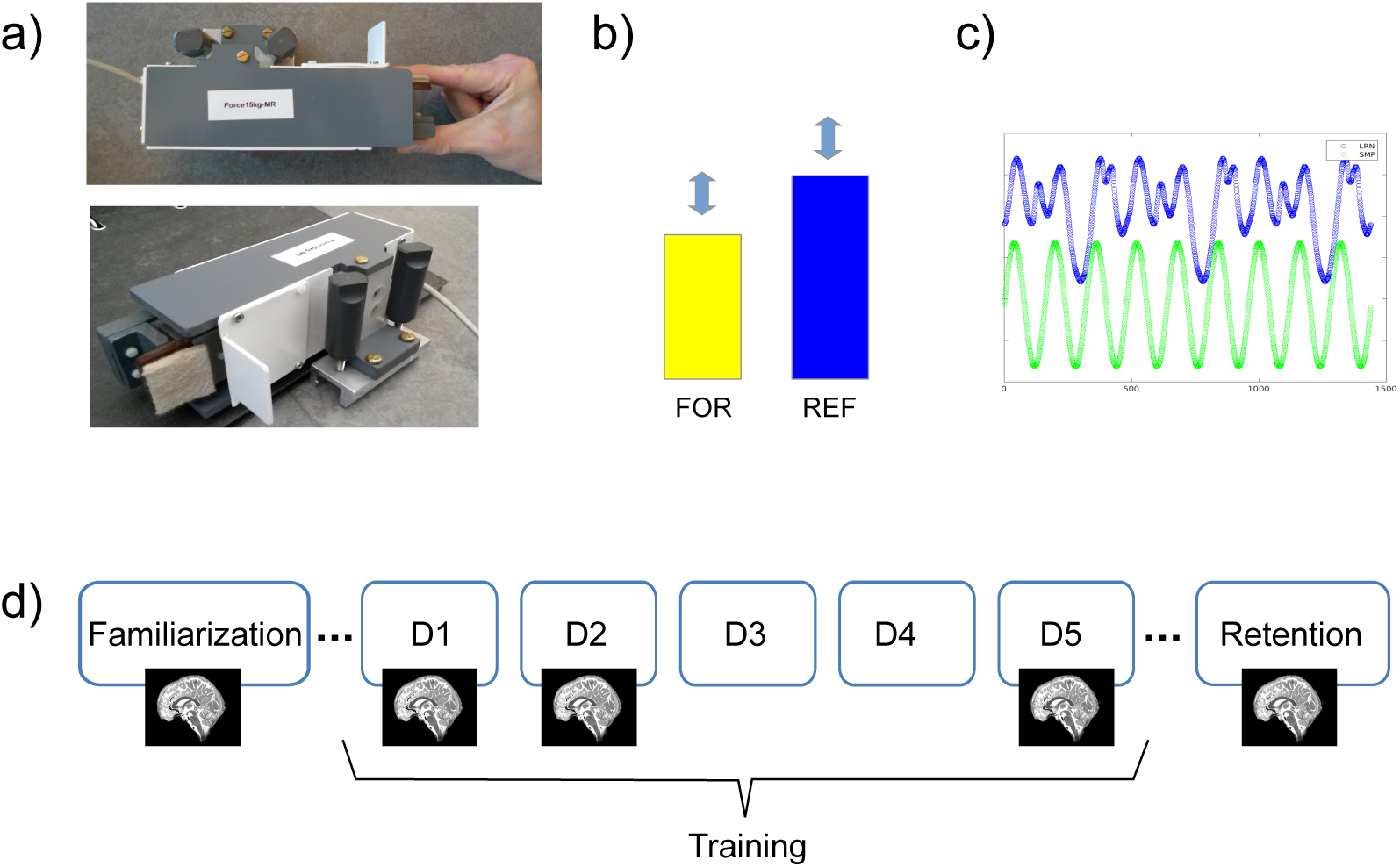
Sequential pinch force task (SPFT) and experimental design. a) SPFT device. Participants hold the pressure sensor between the thumb and index finger. b) They exert force on the sensor to match the height of the reference bar (REF; blue). The FOR bar (yellow) represents the amount of force they are exerting on the device. c) Visual representation of the complex (LRN; blue) and of the simple (SMP; green) sequences. d) Schema of the experimental design. The familiarization session (d0) takes place 2 days before the first day of training (d1) and the retention scan (d17) 12 days after the last training day (d5). All training sessions (d1-d5) take place on consecutive days, Monday to Friday. Participants were in the scanner on d0, d1, d2, d5 and d17.

In the learning condition (LRN), the bar moves following a complex sequence that is difficult to predict (blue on Fig. 1c) (Gryga et al., 2012). In the control condition, the bar moves following a simple sinusoidal sequence (SMP) that is learned almost immediately (green). The SMP sequence was designed to match the LRN sequence in terms of the total magnitude of force, duration, and frequency at the maximum level of force. This control condition can be used to distinguish between potential structural alterations related to motor execution (i.e. participants pinch a device in both conditions) from alterations that are specific to learning a sequence. Lastly, in the rest condition (RST), participants were asked to fixate their gaze on the static REF and FOR bars (both at 50% of their maximal height).

### Experimental Design

The experimental design is illustrated in Fig. 1 (d). Participants performed the SPFT on 5 consecutive days (Monday to Friday). A familiarization session (d0) on the previous Thursday or Friday, 2 days prior to the first day of training, allowed to test the participant’s maximum pinch force and calibrate the level of force required in subsequent training sessions in order to avoid fatigue. The minimum bar level corresponded to a force of 5% of the participant’s maximum force, while the maximum bar level corresponded to 30% of the participant’s maximal force. During this session, participants also became familiar with the device and the task as they performed 9 trials of the SMP sequence. Each training day (d1-d5), participants of the experimental group completed 3 pseudo-randomly presented blocks each consisting of 3 trials of SMP, RST, and LRN, resulting in a total of 9 trials per condition every day. Participants in the control group also performed 3 blocks of training, but LRN trials were replaced by SMP trials. Throughout this manuscript, the experimental group will be referred to as the LRN group and the controls as the SMP group. Each trial lasted 18 seconds and the entire training session lasted 20 minutes as in Gryga et al. (2012). Participants were given feedback on their performance (i.e. average accuracy in matching the height of the REF bar) after the SMP and the LRN trials. A retention session (d17) was conducted approximately 12 days after the last day of training and consisted in the same procedure as the previous training sessions (d1-d5). The task was performed inside the MRI scanner on d0, d1, d2, d5, and d17 and outside the scanner on d3 and d4 (see Fig. 1d). All sessions for all participants took place in the morning to avoid the potential influence of circadian rhythms on our results.

### MRI Acquisitions

MRI data was acquired on a Siemens 7 Tesla scanner (MAGNETOM, Siemens Healthcare, Erlangen, Germany) with a 32-channel Nova head coil at the Max Planck Institute in Leipzig, Germany. DWI data, acquired from an Echo Planar Imaging (EPI) sequence (TR= 10100 ms, TE= 62.8 ms, FOV= 192 x 192 mm^2^, slice acceleration factor: 2, slice thickness = 1.2 mm, 102 slices, GRAPPA factor: 2, partial Fourier 6/8, b=1000 s/mm^2^, 20 directions, PE= AP, bandwidth= 1562 Hz/Px, voxel size= 1.2 ⨯ 1.2 ⨯ 1.2 mm), was used to assess WM microstructure. Rs-fMRI data were acquired with a blood-oxygen-level-dependent (BOLD) sequence (TR= 1130ms, TE= 22ms, flip angle= 40°, FOV = 192 ⨯ 192 mm², slice thickness = 1mm, 102 slices, GRAPPA factor 2, partial Fourier 6/8, bandwidth = 1562 Hz/Px, voxel dimensions = 1.2 ⨯ 1.2 ⨯ 1.2mm). Participants had their eyes open and were fixating their gaze on a cross during this 10-min acquisition. Rs-fMRI and DWI data were acquired prior to SPFT training. Uniform intensity T1-weighted images (UNI) were also acquired with an MP2RAGE sequence (TR = 5000 ms, TE = 2.45 ms, flip angle 1 = 5°, flip angle 2 = 3°, FOV = 224 ⨯ 224 ⨯ 240 mm^3^, slice thickness = 0.7 mm, 240 slices, bandwidth = 250 Hz/Px, voxel size = 0.7 ⨯ 0.7 ⨯ 0.7 mm) (Marques et al., 2010). Fieldmaps were also acquired (TR = 18ms, TE1 = 4.08ms, TE2 = 9.18ms, flip angle = 10°, FOV = 256 ⨯ 256mm², slice thickness = 2mm, 80 slices, bandwidth 1 = 300 Hz/Px, bandwidth 2 = 300 Hz/Px, voxel dimensions = 2 ⨯ 2 ⨯ 2mm) to correct distortions in BOLD images due to field inhomogeneities. As indicated above, one subject from the LRN group was excluded because of a large DWI signal drop in the temporal lobe.

### Image Preprocessing

DWI data were preprocessed using the MRtrix (3.0) software which performs denoising of the data, corrects for motion and Eddy currents (Eddy tool in FSL 6.0.1), and for susceptibility-induced distortions (topup tool in FSL) using b0 volumes of opposing phase-encoding polarities (Andersson et al., 2003; Andersson & Sotiropoulos, 2016; Skare & Bammer, 2010; Smith et al., 2004; Tournier et al., 2019). The gradient scheme containing gradient vectors and b-values (bvecs and bvals) is stored in the header of the MRtrix file format (mif) and automatically reoriented by MRtrix functions. Bvecs and bvals were extracted from the preprocessed image before the next step, which requires the NIFTI format. Bias field correction was performed using the N4 algorithm of ANTs (3.0) within a mask computed using the brain extraction tool (bet) of FSL on the b=0 preprocessed volume (Tustison et al., 2010). A brain extraction of all DWI volumes was then applied using the b=0 mask in order to remove all non-brain voxels. Preprocessed DWI volumes were smoothed anisotropically, a method in which kernels are shaped based on the main directions of fiber tracts, using the 3danisosmooth function in AFNI (19.0.26 ‘Tiberius’) (2 iterations, s1= 0.5, s2= 1.0) (Ding et al., 2005). This type of smoothing has been shown to preserve directional information, maintaining WM structure boundaries and limiting partial voluming effects (Van Hecke et al., 2010). This method was also shown to decrease the influence of smoothing parameters, such as kernel size, on voxel-based analysis results (Jones et al., 2005; Moraschi et al., 2010; Van Hecke et al., 2010).

DWI data were fitted to a tensor model with dwi2tensor (MRtrix 3.0) and the tensor images were converted to a symmetric matrix in the NIFTI format in the lower-triangular ordering (dxx, dxy, dyy, dxz, dyz, dzz) for ANTs. Spatial normalization was performed in ANTs using high-dimensional non-rigid registration of the tensor images, which uses both spatial and directional tensor information, and was shown to improve alignment of WM tracts and minimize shape confounds on FA outcomes (Zhang et al., 2007). Warps were computed at 4 levels: first, using rigid and affine transforms (with mutual information; MI, as similarity metric) to compute the warp from DWI (using the b=0 volume which has the highest contrast) to anatomical space (UNI image from the MP2RAGE T1 acquired in the same session), and then from anatomical (one day) to subject space (across days), from subject to group space, and lastly to MNI space (MNI152) using rigid (MI), affine (MI), and SyN (symmetric normalization; with cross-correlation) transforms with the antsRegistration function (Avants et al., 2008, 2009). The first step of the registration process of tensor images takes care of the spatial alignment: all warps were applied to the tensor images in a single step, using antsApplyTransforms. Linear interpolation in the log space was used and, since the log of 0 is undefined, the background tensor value was set to 0.0007 (Arsigny et al., 2006). All warps were then combined, and the combined transform was used to reorient the deformed tensor images (ReorientTensor), accounting for the orientational aspect of normalization (Zhang et al., 2007). Maps of fractional anisotropy (FA), mean diffusivity (MD), axial diffusivity (AD), and radial diffusivity (RD) were then computed on the reoriented tensor images (in MNI space) using ImageMath in ANTs. A WM mask was created from the mean FA image thresholded at 0.35 to include only WM voxels in statistical analyses.

Rs-fMRI data was corrected for motion and for magnetic field inhomogeneities using the acquired fieldmaps. Nuisance regression, including 12 motion regressors (3 translations and 3 rotations plus their first derivatives), outlier regressors and physiological regressors, was performed using Nilearn’s NiftiMasker. A Gaussian smoothing kernel of 2.4mm was then applied before calculating voxel-wise network centrality metrics degree centrality (DC) and Eigenvector centrality (EC) (Wink et al., 2012). DC and EC maps were non-linearly registered to MNI space with ANTs (Avants et al., 2009). DC and EC provide a measure of the degree of connectivity of a node to other nodes, with each grey matter voxel representing a node. All preprocessing, tissue segmentation and registration scripts, which were implemented in the CBS Tools environment, are openly available at https://github.com/AthSchmidt/MMPI/tree/master/preprocessing.

### Statistical Analyses

#### Performance improvement in the SPFT

Performance and improvements in performance over the course of the training sessions were quantified using a measure of temporal synchronization (SYN) calculated using custom-built scripts in MATLAB (Version R2016a, The MathWorks, Inc., Natick, Massachusetts, United States). SYN was defined as the temporal deviation (in ms) between the time of the movement of the REF bar and the time when the FOR bar matches the height of the REF bar most closely. The time of best match between the REF and FOR patterns was determined using cross-correlation. The time difference (SYN in ms) was then calculated between the time of movement of the REF bar and the time lag with the greatest cross-correlation (i.e., representing the best match between REF and FOR patterns). A SYN score of zero thus indicated perfectly timed performance.

SYN score values of the three trials of each block were averaged for each participant, resulting in three block values per day. Block SYN scores were then averaged, yielding one SYN value per day. A repeated-measures ANOVA was conducted in Jamovi (https://www.jamovi.org; https://cran.r-project.org/; Lenth et al., 2018; Singmann et al., 2018), to assess the progression in performance, with Group (LRN and SMP) as a between-subject factor and Day (1-5) as the repeated-measures factor. Mauchly’s tests were conducted to assess sphericity and the appropriate correction was applied if sphericity was violated (Greenhouse-Geisser if ε < 0.75 or Huynh Feldt if ε > 0.75). Post-hoc Tukey’s tests were then used to assess the specific temporal location of differences in significant effects and interactions (i.e. between which days the improvement in SYN score was significant).

#### WM microstructural changes across learning stages

We conducted voxel-wise analyses within a WM mask on all diffusion maps (FA, MD, AD, and RD) using a flexible factorial design for longitudinal data from the CAT12 (Computational Anatomy Toolbox: http://www.neuro.uni-jena.de/cat/) in SPM12 (Statistical Parametric Mapping software: http://www.fil.ion.ucl.ac.uk/spm/) implemented in MATLAB (Version R2019a, The MathWorks, Inc., Natick, Massachusetts, United States). The flexible factorial design accounts for dependency between time points for each participant, and included two factors: Group (LRN and SMP) and Scan (d1, d2, and d5). Based on the MSL literature, putative changes between d1-d2 were interpreted as occurring during the initial fast learning stage, d2-d5 as the subsequent slow learning stage, and d1-d5 as overall learning. The contrasts assessed were the interaction between Group and Scan in the following manner: d2 > d1, d2 < d1, d5 > d2, d5 < d2, d5 > d1, and d5 < d1. These contrasts were assessed in both groups, only within the LRN group, and with the opposite direction of change in the LRN vs SMP group (e.g. d2 > d1 in LRN, d2 < d1 in SMP), which resulted in a total of 18 contrasts. FWE adjustment was used to correct for multiple comparisons at the cluster-level (p < 0.05). All analyses were conducted within the WM mask. We defined sequence-specific changes as significant changes in the contrasts of opposite directions between groups, where the LRN group was driving the interaction (i.e. greater change in LRN). Significant changes in both groups were defined as non-sequence-specific and interpreted as related to motor execution of the SPFT. Similar voxel-wise interaction analyses were conducted on rs-fMRI centrality metrics with Group and Scan as factors, allowing to identify clusters of sequence-specific changes in functional connectivity (unpublished), which were used for ROI generation.

#### WM microstructural changes associated with sequence-specific functional changes

Work from our group focusing on the rs-fMRI data in the same cohort identified functional reorganization of the networks involved in MSL (unpublished). These regions were used to define ROIs in WM tracts associated with these task-relevant functional changes. Specifically, sequence-specific increases in centrality (DC and EC) were found in the right globus pallidus (GP) in the initial learning stage (d1-d2) and bilaterally in the superior parietal cortex (SPC) in overall learning (d1-d5). Decreases in the right supplementary area (R SMA) and right pars opercularis (R PO) were also observed between d1-d5 in the LRN group. In order to relate structural changes in WM to functional changes, ROIs (Fig. 3, in blue) were generated from the rs-fMRI clusters in grey matter (Fig. 3, in red) using the 3dROIMaker function in AFNI (Taylor & Saad, 2013). The GM ROI was first inflated by two voxels to find where it overlapped WM (within our group WM mask) and then inflated by four voxels to define an ROI within the WM. Inflating parameters were adjusted when creating the R GP and the R SPC ROIs, in order to yield ROIs of similar sizes, as these clusters were located closer to the WM mask. Resulting ROIs contained 79, 187, 238, 161, and 69 voxels for the R GP, L SPC, R SPC, R SMA and R PO, respectively. We extracted mean values from each ROI and conducted repeated-measures ANOVAs across timepoints (d1, d2, and d5) in the LRN group, with separate analyses for each diffusion metric (FA, MD, RD and AD) and ROI. Post-hoc Tukey’s tests were then conducted on significant effects and interactions to determine the locations of significant changes in WM metrics (i.e. between which days). These analyses focused on participants of the LRN group since ROIs from the rs-fMRI data come from an interaction analysis which showed changes specific to the LRN (i.e. sequence-specific) in these regions.R

**Figure 2.**
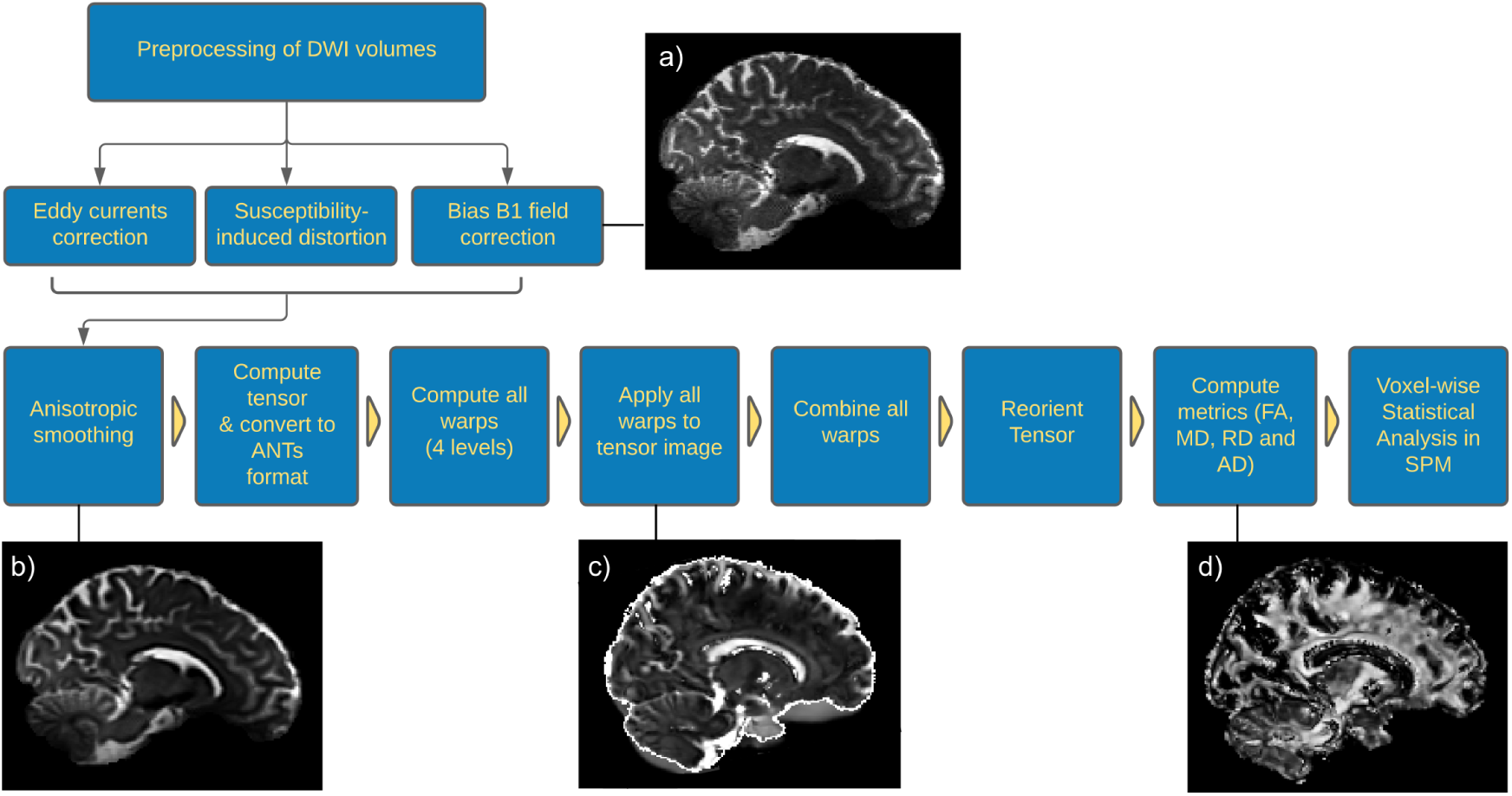
DWI data processing workflow. Preprocessing included correction for motion, Eddy currents, susceptibility-induced distortions, and bias B1 field correction (N4). The preprocessed DWI image (b=0 volume) of one participant is shown in a). Preprocessed DWI volumes were then smoothed anisotropically (b), fitted to a tensor model and converted to a symmetric matrix in the lower-triangular ordering (dxx, dxy, dyy, dxz, dyz, dzz) for ANTs. Four levels of warps were computed (native to T1, T1 to subject, subject to group, and group to MNI) and applied to the tensor image with linear interpolation in the log space in ANTs (c). All warps were combined, and the combined transform was used to reorient the deformed tensor image. Maps of fractional anisotropy (FA), mean diffusivity (MD), axial diffusivity (AD), and radial diffusivity (RD) were computed on the reoriented tensor images. The FA map of the same participant is shown in d. These maps were analyzed with voxel-wise analyses.

**Figure 3.**
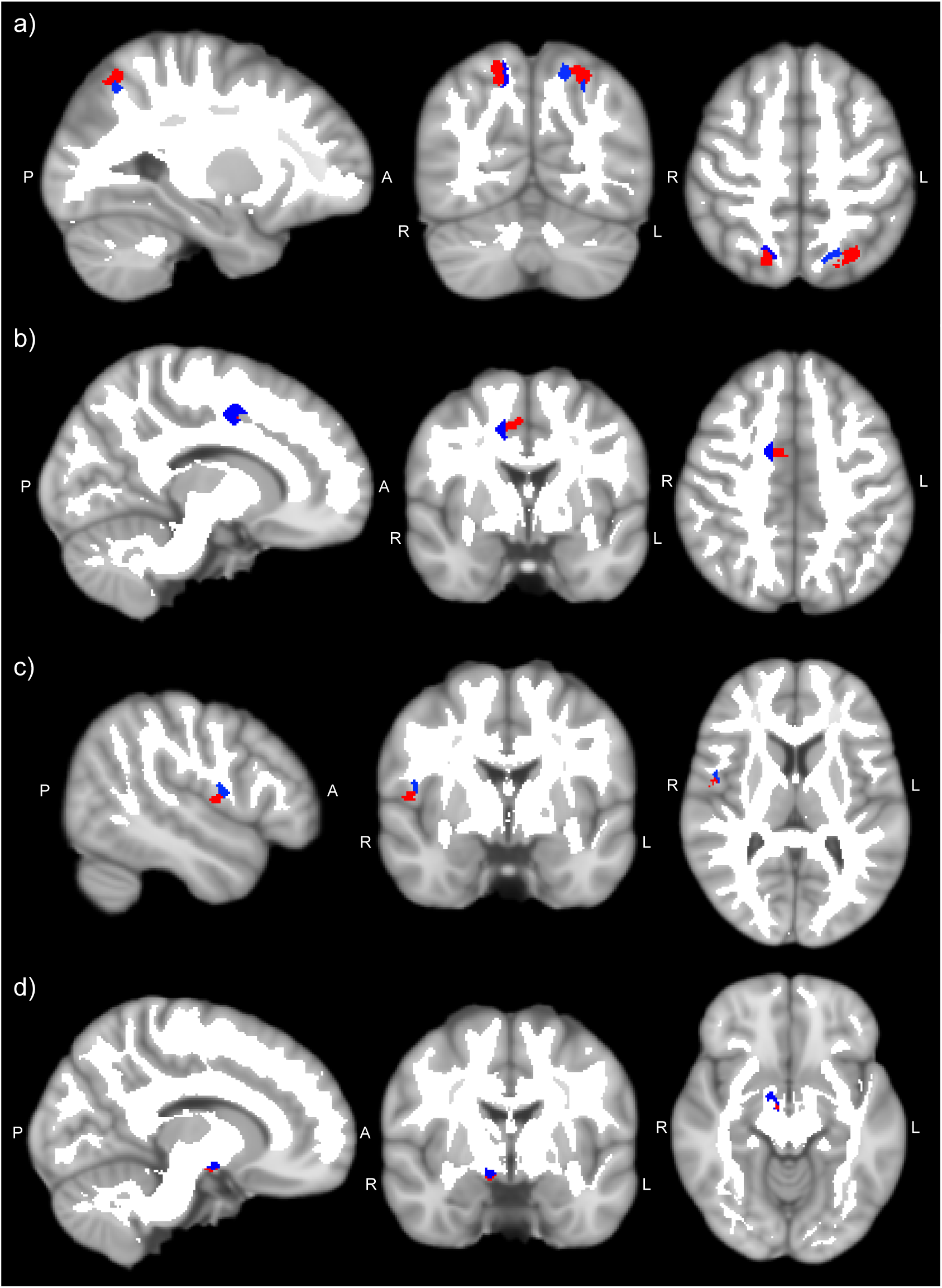
Regions of interest (ROIs) in white matter (blue) created from grey matter ROIs (red) where sequence-specific changes in functional connectivity were found (unpublished). ROIs are displayed on the MNI152 template and WM ROIs are overlaid on a white matter mask created from the group average of FA maps thresholded at 0.35. a) Right and left superior parietal cortex (SPC) ROIs. b) Right supplementary motor area (SMA) ROI. c) Right pars opercularis (PO) ROI. d) Right globus pallidus (GP) ROI.

## RESULTS

### Performance improvement in the SPFT

Participants in the LRN group learned the complex sequence progressively over the course of the training period as evidenced by the large decrease in temporal deviation (SYN) between the beginning of training (d1 block 1 mean SYN score= 224.01 ± 68.53) and the last block of d5 (89.31 ± 62.67). On the other hand, participants in the control group improved very minimally in performing the SMP task as the sequence was fairly easy and temporal deviation was already minimal at the beginning of training (mean SYN score at d1 block 1= 33.98 ± 7.31; mean SYN at last block of d5= 19.23 ± 5.56). Scores were significantly different between groups; a repeated-measures ANOVA revealed a significant main effect of Group (F(1,35)= 84.0, p < 0.001) and an effect size (η^2^ _p_) of 0.706, further showing how different the sequences are (see Fig. 4). There were no significant differences in performance improvement between musically active participants and those who were not (p > .05) and the level of physical exercise showed no significant correlation with performance improvement (p > .05).

**Figure 4.**
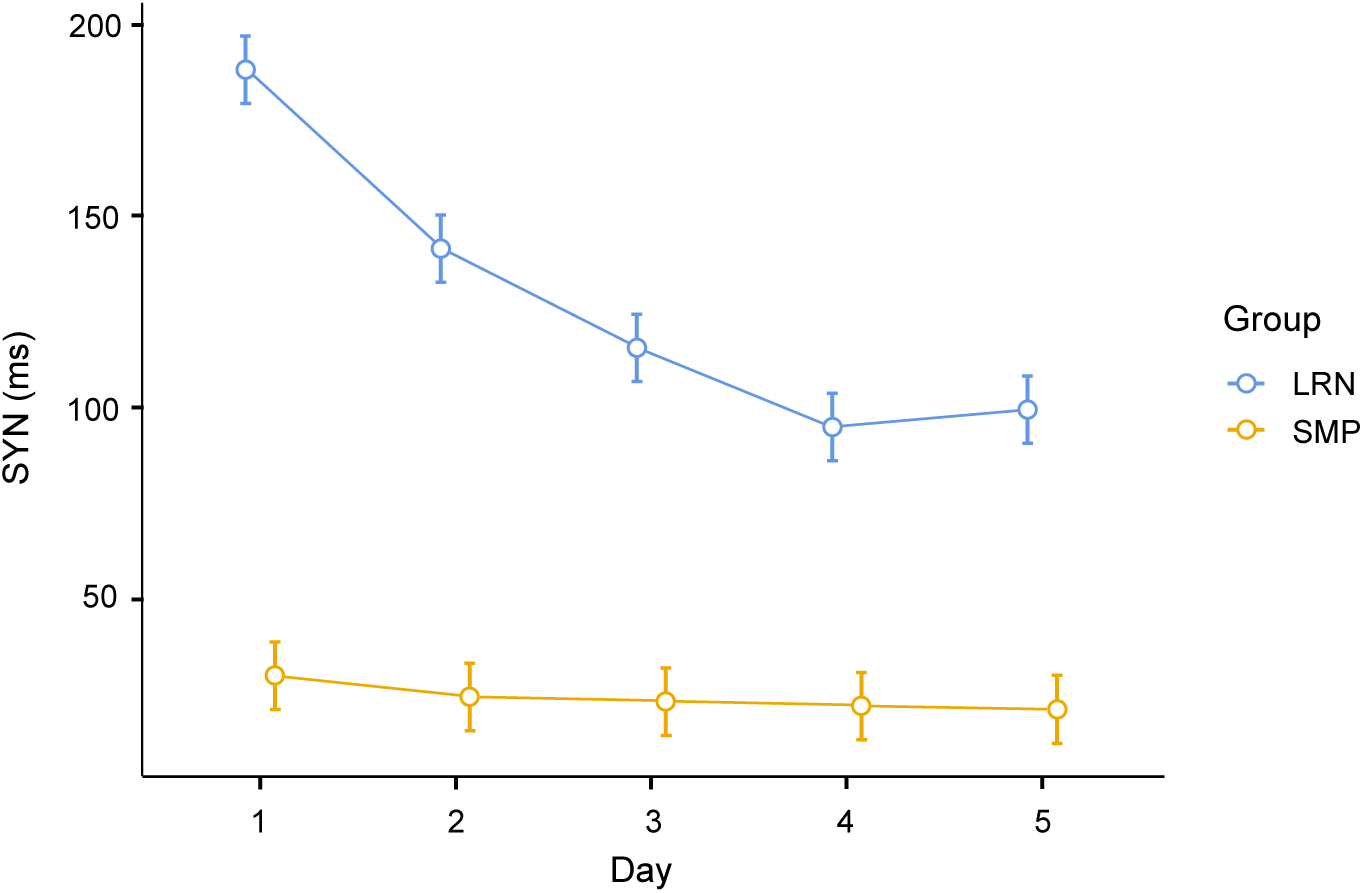
Behavioral Results. Temporal deviation (SYN; in ms) for both groups across training days (d1- d5), where the SYN value of each day is the mean across blocks. LRN: learning group (in blue); SMP: control group (in orange). Error bars represent the standard error of the mean.

There was also a main effect of Day (F(4,140)= 52.7, p<0.001, η^2^ _p_= 0.601) and a significant interaction of Day*Group (F(4,140)= 37.0, p<0.001, η^2^ _p_= 0.514). These main effects and interactions were still significant (p<0.001) after Greenhouse-Geisser correction which was applied because sphericity was violated in this analysis. Consistent with the theories of learning stages, most improvements took place in the first days and then reached a plateau at d4 in the LRN group; post-hoc Tukey was significant between d1-d2 (t=8.235, p<0.001), d2-d3 (t= 4.573, p<0.001), and d3-d4 (t= 3.662, p=0.013), but not significant between d4-d5 (t= -0.810, p= 0.998) (Fig. 4).

In contrast, participants in the SMP group exhibited little significant improvement over the course of the 5 days of training. None of the pairwise comparisons between consecutive days were significant in this group and Post-hoc Tukey was also non-significant between d1-d5, indicating no significant improvement in the overall learning period. Two participants from the SMP group were excluded from the analysis because their SYN scores on d3, and d4 for one of them, were outliers as they exceeded the mean by more than two SD.

### WM microstructural changes across learning stages

Changes of opposite directions in both groups were found in the corticospinal tract, underlying the left primary motor cortex (M1) (T= 4.20, p= 0.002; Fig. 5b). FA decreased in LRN, whereas FA increased in this cluster in SMP (d5 < d1 in LRN, d5 > d1 in SMP). The mean ΔFA in LRN was - 0.029, while FA increased by 0.0395 in the SMP group. Voxel-wise analyses also revealed a decrease in FA (T= 5.01, p= 0.005; mean ΔFA= -0.176) in the right ascending sensorimotor tract adjacent to the primary somatosensory cortex (S1), in the LRN group only, during overall learning (d5 < d1; see Fig. 5a). Other DTI metrics for these contrasts were non-significant (p_FWE_ > 0.05).

**Figure 5.**
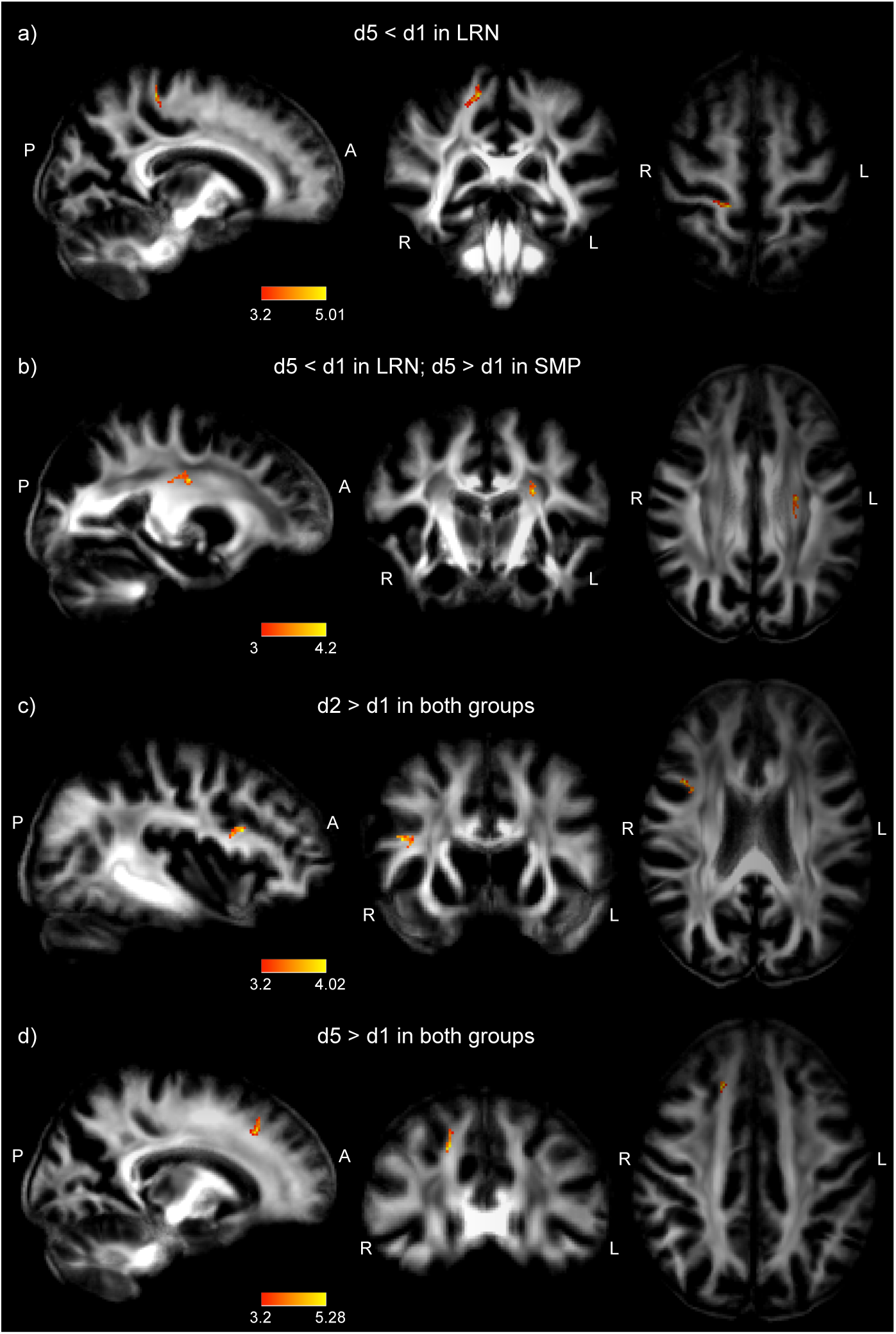
Changes in FA from voxel-wise analyses. T-stat maps (maximum intensity projection for better visualization) are overlaid on the mean FA image. a) Decrease in FA in the LRN group between d1-d5 in the right ascending sensorimotor tract connecting to the primary somatosensory cortex (S1). b) Decrease in FA in LRN and increase in FA in SMP between d1-d5 in the left corticospinal tract connecting to the primary motor cortex (M1). c) Increase in FA in both groups (related to motor execution) between d1-d2 in the right frontal inferior longitudinal (FIL) tract connecting to the pars opercularis (PO). d) Increase in FA in both groups (related to motor execution) between d1-d5 in anterior corona radiata connecting to the right frontal eye field (FEF).

There were also changes in FA in the same regions in both groups. These plastic changes were common to both groups and therefore considered non-sequence specific and more related to motor execution. FA had a near significant increase in the frontal inferior longitudinal (FIL) tract underlying the right pars opercularis (PO) in the early stage of learning (d2 < d1; Fig. 5c) in both groups (T= 4.02, p= 0.063; mean ΔFA in all subjects= 0.115, mean ΔFA in LRN= 0.125, mean ΔFA in SMP= 0.104). In overall learning, FA increased significantly in the right anterior corona radiata adjacent to the frontal eye field (FEF; T= 5.28, p=0.023; Fig. 5d) in both groups (mean ΔFA in all subjects= 0.119, mean ΔFA in LRN= 0.147, mean ΔFA in SMP= 0.093). Other metrics for these contrasts, and all other contrasts assessed, were non-significant (p_FWE_ > 0.05). The results are summarized in Table 2.

**Table 2.**
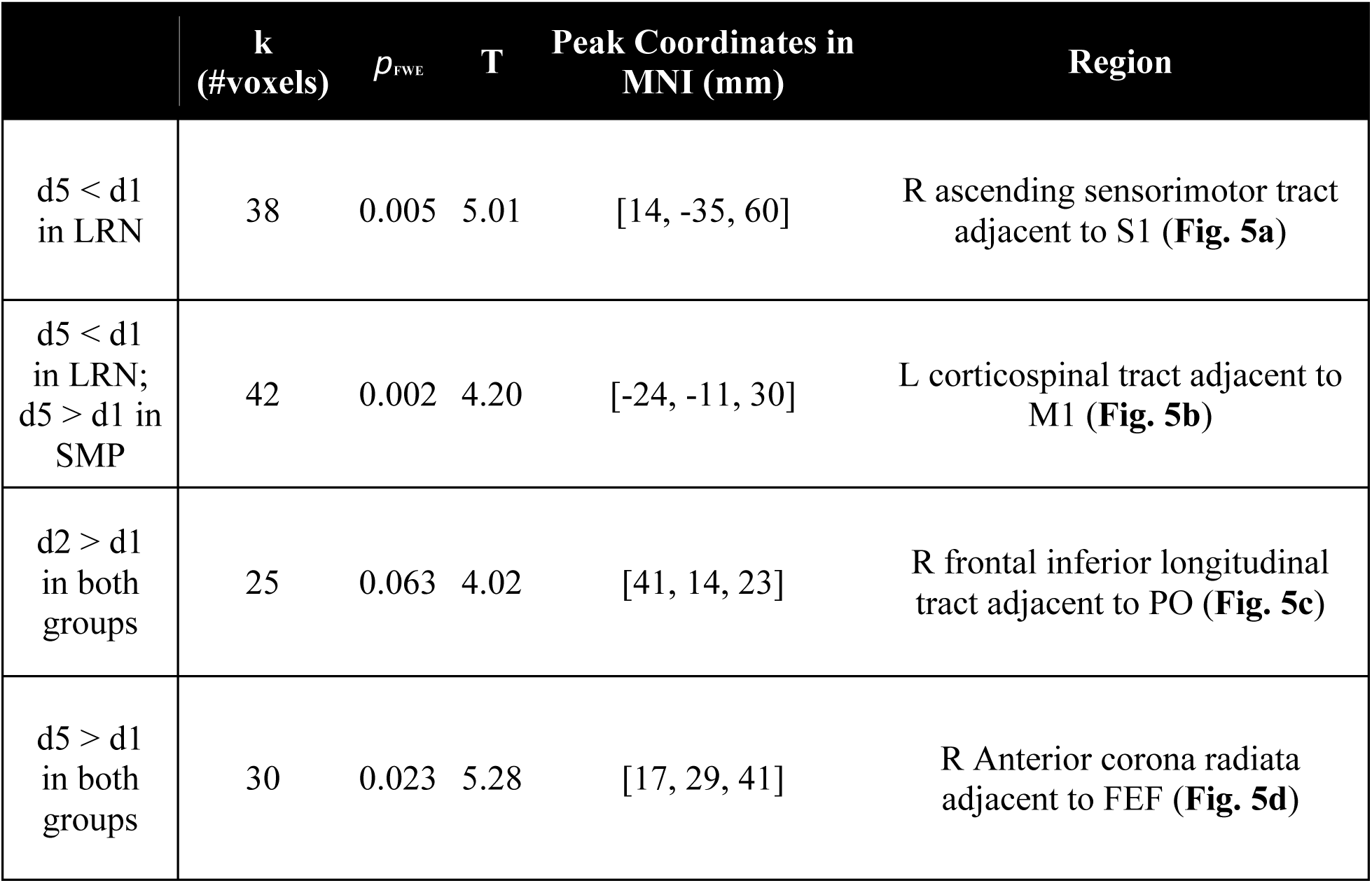
Clusters in which significant changes in FA were found.

|  | **k**<br>**(#voxels)** | **$p_{\text{FWE}}$** | **T** | **Peak Coordinates in**<br>**MNI (mm)** | **Region** |
| --- | --- | --- | --- | --- | --- |
| d5 < d1<br>in LRN | 38 | 0.005 | 5.01 | [14, -35, 60] | R ascending sensorimotor tract<br>adjacent to S1 (**Fig. 5a**) |
| d5 < d1<br>in LRN;<br>d5 > d1 in<br>SMP | 42 | 0.002 | 4.20 | [-24, -11, 30] | L corticospinal tract adjacent to<br>M1 (**Fig. 5b**) |
| d2 > d1<br>in both<br>groups | 25 | 0.063 | 4.02 | [41, 14, 23] | R frontal inferior longitudinal<br>tract adjacent to PO (**Fig. 5c**) |
| d5 > d1<br>in both<br>groups | 30 | 0.023 | 5.28 | [17, 29, 41] | R Anterior corona radiata<br>adjacent to FEF (**Fig. 5d**) |

### WM microstructural changes associated with functional changes

WM microstructure in the ROI underlying the right SMA was altered during the training period in the LRN group (Fig. 6a). FA was found to decrease significantly across days (F(2, 36)= 5.82, p=0.006, η^2^ _p_= 0.244; Fig. 6b) and Tukey’s post-hoc test revealed that this decrease was significant between d1-d2 (t= 3.072, p=0.011) and between d1-d5 (t= 2.823, p=0.021), but not between d2-d5 (t= -0.249, p=0.966). There was also a significant decrease in AD (Fig. 6c) in the R SMA (F(1.38, 24.91)= 6.27, p=0.012 after Greenhouse-Geisser correction, η^2^ _p_= 0.258) and post-hoc tests showed that the differences were statistically significant between d1-d2 (t= 3.300, p= 0.006) and between d1-d5 (t= 2.763, p= 0.024). RD increased significantly across days (F(1.44, 25.99)= 3.93, p= 0.044 after Greenhouse-Geisser correction, η^2^_p_= 0.179) (Fig. 6d). This increase bordered on statistical significance in Post-hoc Tukey’s tests between d1-d2 (t= -2.419, p=0.053) and between d1-d5 (t= -2.438, p= 0.051), but not between d2-d5 (t= -0.0190, p= 1.000). MD showed no significant change in this ROI (p > 0.05).

**Figure 6.**
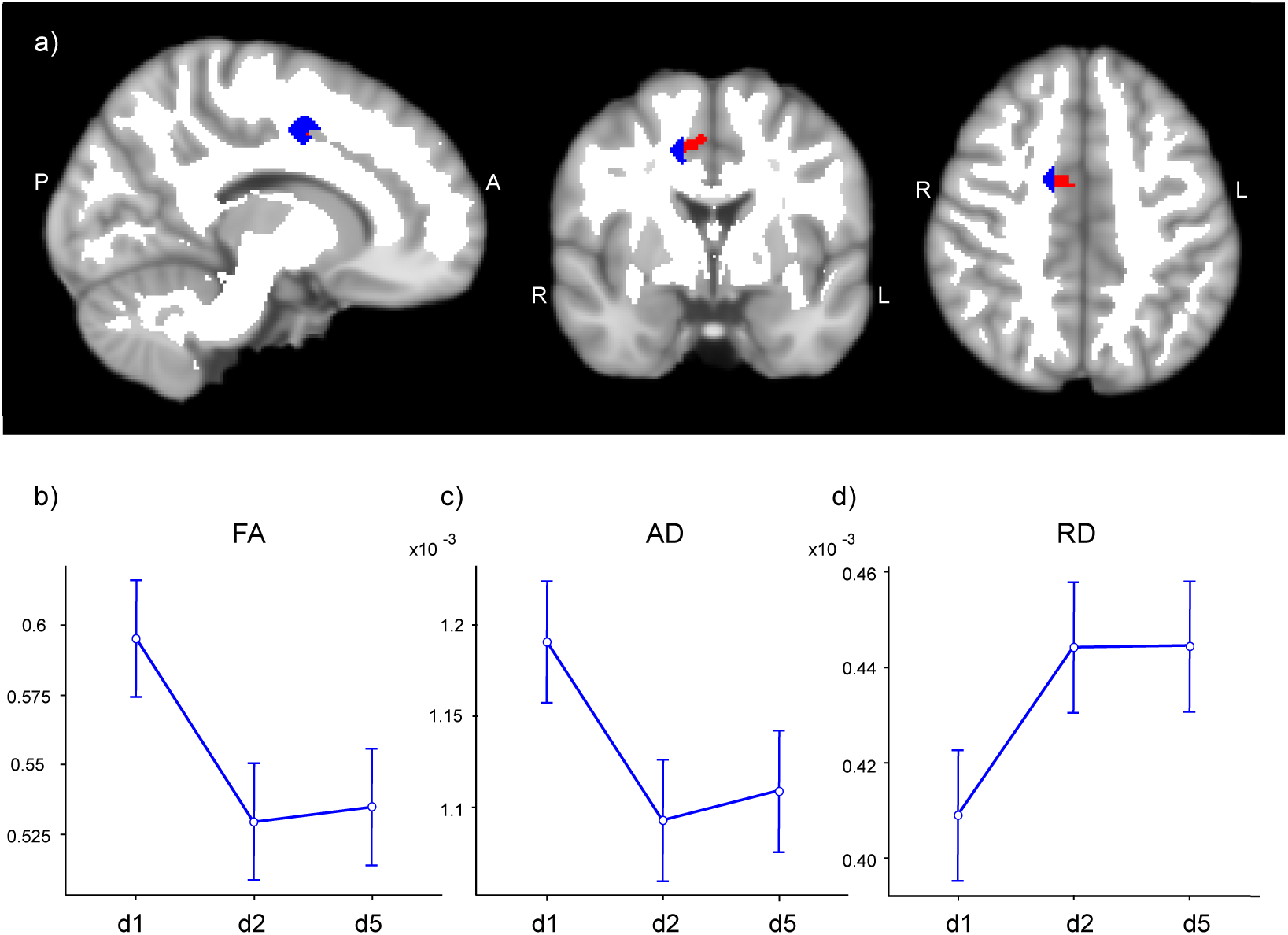
Changes in WM microstructure in the ROI underlying the right supplementary area (SMA) in which sequence-specific changes in functional connectivity were found (unpublished). a) The right SMA ROI from resting-state analyses (in red) and the WM ROI (in blue; overlaid on the WM mask in white) are both overlaid on the MNI152 template. b-c) FA and AD decreased in the LRN group between d1 and d2 and remained lower at d5. d) RD increased between d1 and d2 in LRN and remained higher at d5.

Diffusion metrics in the other ROIs (L and R SPC, R GP, and R PO) did not change significantly across days in repeated-measures ANOVA analyses.

## DISCUSSION

In this study, we investigated structural changes in WM over the course of 5 training days on a continuous visuomotor sequence task using DTI. Consistent with the behavioral results, where the greatest amount of improvements in temporal synchronization (SYN) were detected between the two first days (see Fig. 4), we observed structural changes in WM only in the early phase of learning (d1-d2), and when looking at the overall learning period (d1-d5), which suggests a slower, more progressive change. Sequence-specificity was assessed through interaction analyses between the LRN group, who performed a complex sequence, and the control group (SMP), who performed a simple sequence, where the interaction is driven by the LRNs. FA was found to change in opposite directions in both groups in the left corticospinal tract (CST) inferior to the primary motor cortex (M1; Fig. 5b) during overall learning. However, as the SMP group showed a greater change than the LRNs, we cannot establish that altered FA in WM underlying M1 is due to sequence learning per se. Changes in the right ascending sensorimotor tract (SMT) adjacent to the primary somatosensory cortex (S1) were also observed in the LRN group (Fig. 5a). WM microstructure was altered during the early phase of learning in the ROI underlying the right supplementary motor area (SMA; Fig. 6), where sequence-specific changes in functional connectivity were found in this cohort (unpublished). Together, our findings provide evidence for training-dependent white matter plasticity in the sensorimotor network during short-term motor sequence learning.

### Changes in the LRN group

#### Overall Learning - Right Primary somatosensory cortex (S1)

Fractional anisotropy decreased in the right ascending SMT in participants of the LRN group during overall learning (d1-d5; Fig. 5a). We hypothesize that this decrease in FA in fiber tracts connecting to the right S1 may reflect suppression of activity in S1 ipsilateral to the hand used in the SPFT (Kastrup et al., 2008; Staines et al., 2002). Increased activity in the contralateral S1, and suppression of activity in the ipsilateral S1, as a result of task-relevant somatosensory stimulation and voluntary movements, have been reported in fMRI and EEG studies (Kastrup et al., 2008; Lei & Perez, 2017; Nirkko et al., 2001; Staines et al., 2002). Blood flow suppression along with inhibition of S1 areas that are not involved in a task (e.g. ipsilateral body parts) from the prefrontal-thalamic system have been suggested as mechanisms to selectively gate sensory inputs (Drevets et al., 1995; Knight et al., 1999; Staines et al., 2002; Yamaguchi & Knight, 1990). In work by Drevets and colleagues (1995), blood flow reductions were observed in S1 ipsilateral to the expected stimulus when participants were anticipating somatosensory stimulation. The fact that changes in the SMT were only found in the LRN group may be due to increased levels of anticipation and attention in participants performing a complex task (Drevets et al., 1995; Halsband & Lange, 2006; Staines et al., 2002). We speculated that the increased attention necessary to perform the complex task would likely require greater activation of the dorsolateral prefrontal cortex. This could lead to greater inhibition of the ipsilateral S1, through the prefrontal-thalamic sensory gating system, as a way to suppress background noise and enhance processing of task-relevant inputs (Corbetta & Shulman, 2002; Halsband & Lange, 2006; Staines et al., 2002; Yamaguchi & Knight, 1990). Participants of the SMP group on the other hand may not need to pay as much attention to the task at hand, and to the associated sensory inputs, and may perform the repetitive sinusoidal sequence in a more automated manner.

This sensory gating, in which behaviorally-irrelevant regions are inhibited in order to effectively suppress unimportant, and potentially disruptive, inputs, may thus contribute to enhancing the responsiveness of the contralateral S1 to stimuli (Staines et al., 2002). Enhanced somatosensory inputs while performing a motor task have been associated with improvements in performance, and disrupting somatosensation during training impairs motor learning (Vidoni et al., 2010; Wei et al., 2018). As participants in this study were holding a pressure sensor between their thumb and index fingers, enhanced somatosensory inputs to the central nervous system due to SPFT training could result in the acquisition of a new task-specific sensory map (Braun et al., 2000; Pascual-Leone et al., 2005). This would provide better sensory feedback when subsequently performing the task, which could translate into improved accuracy in matching the reference bar (Wei et al., 2018).

#### Overall Learning - Left Primary motor cortex (M1)

FA in the left CST connecting to M1 was found to decrease in the LRN group, while it increased in the SMP group during overall learning (d1-d5; Fig. 5b), indicating a relatively slow change in this region over the course of the five training days. As the SMP group did not show any significant improvement in the SPFT, we are cautious in interpreting a change in this group without supporting behavioral evidence. The change in the LRN group on the other hand was accompanied by a change in behavior. We will thus focus the interpretation on the LRN group although we could speculate that any plastic change occurring in the SMP group could mirror changes in the LRNs but occur on a shorter time scale (Dayan & Cohen, 2011; Hyde et al., 2009; A. Karni et al., 1995).

Since we expect M1 to be activated when performing a motor task with the contralateral limb, as M1 has a known role in motor execution and the storage of learned sequence representations (Bengtsson et al., 2005; Hardwick et al., 2013; Hyde et al., 2009; Monfils et al., 2005; Penhune & Steele, 2012; Yokoi et al., 2018), this decrease in FA in the group performing a more complex task may seem contradictory. However, activity in M1 was previously shown to progressively decrease as a motor skill is learned (Dayan & Cohen, 2011; Poldrack, 2000; Seidler et al., 2005), possibly reflecting increased efficiency. Moreover, functional connectivity in M1 was found to increase in the LRN group during the early stage of training (unpublished). This suggests an important and active role of this area as we begin to learn a task, but then, as the motor sequence is learned, less neuronal resources would be needed to perform the task which could be reflected by decreased connectivity (Dayan & Cohen, 2011; Poldrack, 2000). In line with this, studies in musicians, experts in sensorimotor control, have also reported lower FA in motor circuits, including the bilateral CST and corona radiata (Imfeld et al., 2009; Penhune & Steele, 2012 for review; Schmithorst & Wilke, 2002). This decrease in anisotropy may be due to increased efficiency, or it may result from changes in the permeability of axonal membranes to water, or to an increased axonal diameter, which would lead to an increase in intracellular radial diffusivity (Beaulieu, 2002; Imfeld et al., 2009; Zatorre et al., 2012). Lastly, the development of a secondary fiber population in areas of crossing fibers is another potential mechanism through which FA could be reduced (Zatorre et al., 2012). Indeed, another study investigating structural changes associated with five days of MSL found that lower FA in the CST on the last training day correlated with better performance on the task (Steele et al., 2012). Since the significant correlation was located in a region where the CST and superior longitudinal fasciculus (SLF) cross, they hypothesized that maturation of the SLF, which would be the secondary fiber population here, drove the reduction in FA and promoted performance, as the SLF connects cortical regions that are involved in this task. The lack of significant changes in other diffusivity metrics in this study however does not allow to disentangle the extent to which these factors contribute to the reduction in FA observed in the CST. Acquiring a greater number of diffusion gradient directions and strengths, and using more advanced diffusion models such as neurite orientation dispersion and density imaging (NODDI), and tractography, may allow to distinguish the underlying factors leading to altered FA in future studies (Steele & Zatorre, 2018; Tardif et al., 2016; Zhang et al., 2012). However, the increase in functional connectivity observed during the early stage of learning (unpublished), along with the slower decrease in FA found in this study, support the hypothesis of a strong initial M1 involvement, followed by decreased M1 activation as learning progresses, reflecting enhanced network efficiency (Bassett et al., 2015; Costa et al., 2004; Dayan & Cohen, 2011; Finc et al., 2020; A. Karni et al., 1995; Mohr et al., 2016; Poldrack, 2000).

#### Fast Learning - Right Supplementary motor area (SMA)

ROI-based analyses were conducted in WM tracts underlying clusters of sequence-specific changes in resting-state centrality metrics, which included bilateral SPC, right GP, right SMA and right PO (inferior part) (Fig. 3). These analyses revealed changes in the WM ROI underlying the right SMA, in a fiber pathway we identified as the frontal aslant tract (FAT) (Fig. 6a). The FAT connects the superior frontal gyrus (SFG), including the supplementary motor area (SMA), to the IFG (Briggs et al., 2018) and is thought to have a role in working memory, motor planning and coordination (Dick et al., 2019; Varriano et al., 2020). This pathway may function to coordinate sequential motor movements, especially in visuo-spatial tasks, to select the appropriate motor outputs (Dick et al., 2019). Moreover, the SMA has been shown to be involved in long-term (Hikosaka et al., 1995, 1996; Jenkins et al., 1994; Shima & Tanji, 2000; Tanji, 1996; van Mier et al., 1998), and short-term MSL (Vollmann et al., 2013), especially in action planning and in the organization of temporal aspects, such as timing and order, in a wide range of domains (e.g. language, working memory, motor sequences) (Cona & Semenza, 2017). A study using the same task provided strong evidence of the implication of the SMA in sequence learning by showing that non-invasive stimulation over the SMA led to improvements in performance of the SPFT (Vollmann et al., 2013).

Considering the known role of the SMA and FAT in sequence processing (Cona & Semenza, 2017; Dick et al., 2019; Shima & Tanji, 2000; Tanji, 1996; Varriano et al., 2020; Vollmann et al., 2013), we may expect connectivity to be enhanced with this type of task. However, work from our group has found decreases in functional connectivity between the right SMA and right PO during overall learning in the same cohort (unpublished). It was hypothesized that reduced connectivity between these regions may thus reflect increased efficiency, or decreased need for resources to plan and coordinate movements, which would result in segregation of the network. This hypothesis is consistent with the view that the integration of multiple large-scale networks is necessary in the early stages of learning (Finc et al., 2020) and then, as a skill is mastered and becomes nearly automatic, a more easily reachable network, consisting in autonomous segregated modules, is sufficient to execute the task (Bassett et al., 2015; Finc et al., 2020; Mohr et al., 2016). In the present study, we found decreases in FA and AD in the WM ROI underlying the right SMA in the LRN group, during the early stage of learning and in overall learning (Fig. 6a-c). These findings, along with the increase in RD during the same period (Fig. 6d), suggest a decrease in structural connectivity. This is in line with the reduction in functional connectivity found in the right SMA (unpublished).

The spatial and temporal patterns of WM structural changes observed in the current study are in line with a theoretical framework describing two networks operating in parallel during MSL with different time courses (Hikosaka et al., 2002). Those two networks would each subserve distinct aspects of MSL: learning spatial coordinates, supported by a prefronto-parietal loop, and learning motor coordinates, which occurs more slowly and is supported by a M1-sensorimotor loop (Dayan & Cohen, 2011; Hikosaka et al., 2002). Both of these loops also receive contributions from different parts of the striatum and cerebellum depending on the learning stage and on the component learned (spatial vs motor) (Hikosaka et al., 2002; Penhune & Steele, 2012). Interestingly, we observed changes only in the motor loop (M1 and S1). This may be due to the fact that learning spatial coordinates requires less time than learning motor coordinates (Hikosaka et al., 2002; Miller & Cohen, 2001; Penhune & Steele, 2012). This could mean, in the context of the SPFT, that spatial learning took place within the first session, perhaps on the timescale of minutes or hours considering the low complexity of the spatial coordinates to be learned in this task, and did not induce changes in WM that we could detect considering the much longer timescale of our measurements (i.e. days). Moreover, as a sequence is learned, its performance becomes more implicit, and thus relies more heavily on motor mechanisms and very little on attention-demanding spatial mechanisms (Hikosaka et al., 2002; Penhune & Steele, 2012). The SMA, another area in which we observed changes, both functionally (unpublished) and structurally, would provide the link between those two parallel loops, allowing for updated spatial representations to be used by the M1-sensorimotor loop to optimize motor output, according to this framework (Hikosaka et al., 2002). Indeed, Penhune & Steele (2012) emphasized the need for a high degree of interaction between these parallel systems to optimize MSL. The time course of changes in WM tracts underlying the SMA (fast change; d1-d2), and S1-M1 (slower changes; d1- d5), is also in line with the hypothesis that the SMA is involved in converting quickly acquired spatial coordinates to motor coordinates which are then processed by the M1-sensorimotor loop (Hikosaka et al., 2002).

Our findings are in line with previous literature (Bloechle et al., 2016; A. Karni et al., 1995; Klein et al., 2019; Lotze et al., 2003; Schmithorst & Wilke, 2002; Zatorre et al., 2012) and point to the highly dynamic plastic processes in WM tracts underlying the SMA-M1-sensorimotor loop, which parallel functional changes. It has been suggested that increases in anisotropy may reflect ongoing enhancement of fibers organization while decreased FA may be related to increased network efficiency in later stages of learning (Schmithorst & Wilke, 2002).

### Changes in both groups

Changes in anisotropy were also observed in both groups, suggesting the involvement of a number of regions in motor execution rather than sequence-specific learning. FA increased in WM tracts underlying the inferior frontal gyrus (IFG; opercular part) in the early stage of training (d1-d2; Fig. 5c), and in the right anterior corona radiata (aCR) adjacent to the right frontal eye field (FEF) during overall learning (d1-d5; Fig. 5d).

#### Overall Learning - Right Frontal Eye Field (FEF)

The frontal eye field (FEF) is involved in processing visual inputs and controlling voluntary eye movements and its activation is thought to be dependent on the saliency of the target (i.e. whether the target is behaviorally relevant) (Schall & Bichot, 1998; Vernet et al., 2014). In this task, participants maintained the gaze on the computer screen where the REF and FOR bars (Fig. 1) provided both instructions (REF) and feedback (FOR) for the SPFT. Maintaining the gaze on a visual target for extended periods of time (∼20 min/day for 5 consecutive days) may require high sustained activation in the FEF which could translate into structural changes in fiber tracts connecting this region during the overall learning period. The FEF clearly plays an important role in visually guided tasks, but its role has been investigated mostly in the context of goal-oriented saccadic eye movements (Schall & Bichot, 1998). However, in addition to its role in target selection in saccades, the FEF is also involved in the detection and analysis of visual inputs during periods of fixation of the gaze (Posner, 1980; Schall, 2004; Schall & Bichot, 1998). Non-invasive neurostimulation of the right FEF was shown to enhance visual perception and improve performance in a visual detection task (Chanes et al., 2012). Other studies support the idea that the FEF, especially of the right hemisphere, is involved in shifting visual attention without eye movement (Donner et al., 2000; Grosbras & Paus, 2002). Moreover, the FEF may play a role in short-term memory of visuo-spatial information (Clark et al., 2012; Gaymard et al., 1999). The high potential for plasticity of the FEF has made this region a target for neurostimulation to increase visuo-spatial attention in healthy and patient populations (Vernet et al., 2014). Our results, showing FA increases in both groups, support the view that the FEF is highly plastic and suggest that this region is relevant in directing visual attention regardless of task complexity.

#### Fast Learning - Right Pars Opercularis (PO)

Increased FA was also observed in the frontal inferior longitudinal (FIL) tract underlying the dorsal part of the right pars opercularis (PO) between d1-d2 (Fig. 5c). The FIL tract is a chain of u-shaped fibers connecting the dorsal part of the IFG (including the PO), to the middle frontal gyrus and pre-central gyrus (M1) (Catani et al., 2012). U-fibers of the frontal and parietal lobes have been shown to play an important role in sensorimotor integration (Catani et al., 2012, 2017), and have been hypothesized to coordinate movement planning and execution by linking motor and premotor regions (Catani et al., 2012). Moreover, the right PO has been specifically linked with fine motor control of manual motion (Briggs et al., 2019; Liakakis et al., 2011).

Our findings suggest that high task complexity might not be necessary to recruit the PO and incur structural changes in the underlying WM tracts, as the SMP sequence also requires the integration of sensory information to execute the task, as well as fine motor control in order for the appropriate amount of force to be applied on the device at the right time. Furthermore, the change in fiber tracts underlying the PO was observed in the early stage of learning, which may indicate a greater need for sensorimotor integration at this stage.

### Limitations and Future Considerations

The main limitation of this study is that the high field strength (7T) may make our findings less generalizable across studies, as 3T is still much more commonly used in research. Moreover, despite the high spatial resolution of our acquisition, the angular resolution was low, with only 20 directions, and a single diffusion gradient strength was applied (i.e. one shell). The angular sampling, number of diffusion shells and gradient strength were limited due to time constraints, as the study involved the acquisition of several other MRI sequences of long duration. In future studies, DWI acquisitions with higher gradient strengths and a greater amount of directions would allow for tractography to be performed, which would increase certainty when identifying fiber tracts where changes in scalar DTI metrics are observed, and for tract-based quantification of DTI metrics (Mukherjee et al., 2008; Wakana et al., 2007). Furthermore, with a higher number of shells and directions, more advanced modelling approaches, such as NODDI, can be used, which would allow to disentangle factors such as fiber density and orientation dispersion, especially in areas of crossing fibers (Steele & Zatorre, 2018; Tardif et al., 2016; Zhang et al., 2012).

Another challenge when investigating neuroplasticity in MSL is that high variability in the duration of each learning stage, depending on the complexity of the task, makes it difficult to relate stage-specific findings across studies (Dayan & Cohen, 2011; Hyde et al., 2009; A. Karni et al., 1995). Moreover, our experimental design did not allow us to distinguish structural changes occurring during consolidation (i.e. offline), from those occurring during training (i.e. online). However, it is unlikely that the techniques used in the current study would have had the sensitivity to detect such subtle differences and the structural changes observed in WM are likely the sum of alterations taking place both during the training session (i.e. online) and in between sessions (i.e. offline).

Lastly, the motor sequence training period was of short duration in the present study which may limit the amount of observable structural changes. A longer training duration may have led to a greater amount of plastic changes in WM tracts which could have provided further insights into MSL-related neuroplasticity.

## CONCLUSION

Our study provided evidence for white matter plasticity in the sensorimotor network, where the SMA plays a role in linking the spatial and motor aspects, in short-term learning of motor sequences. Our findings also highlighted the time course of plastic changes in this network as we scanned participants not only in the beginning and at the end of training, but also on the second day, allowing for the characterization of changes occurring in the early stage of training. Future ultra-high field MRI studies investigating plasticity in the context of MSL should use a high angular resolution, and a higher number of diffusion shells of varying strengths. This would provide more precision in localizing areas of change and in characterizing the biological underpinnings of plastic changes in brain white matter.

## Acknowledgments

We would like to thank Elisabeth Wladimirov and Domenica Wilfling for their help and involvement in data acquisition and logistics of the multi-modal plasticity initiative (MMPI) dataset. This work was supported by the Max Planck Society, the MaxNetAging Research School (Max Planck Institute for Demographic Research, ATJ), the NWO Vici grant (PI: Birte Forstmann)(PLB), the National Science and Engineering Research Council (NSERC; CJG: RGPIN 2015-04665, CJS: RGPIN-2020-06812, DGECR-2020-00146), the Michal and Renata Hornstein Chair in Cardiovascular Imaging (CJG), the Heart and Stroke Foundation (New Investigator Award, CJG and CJS), the Canadian Institutes of Health Research (HNC 170723), the Fonds de Recherche du Québec - Nature et Technologies (CJS and JH), the Fonds de Recherche du Québec - Santé (JH), and the Réseau de Recherche en Santé cardiométabolique, diabète et obésité (CMDO; JH).

## Authors’ contributions

All authors contributed to the study and approved the final manuscript. Detailed contributions, described according to CRediT, were as follow: Conceptualization: Christine L. Tardif, Pierre-Louis Bazin, Christopher J. Steele, and Claudine J. Gauthier; Methodology: Christine L. Tardif, Pierre-Louis Bazin, and Christopher J. Steele; Data collection: Uta Schneider, Sophia Grahl, Anna-Thekla Jäger and Julia Huck; Formal analysis: Stéfanie A. Tremblay, Anna-Thekla Jäger, Chiara Giacosa, and Stephanie Beram; Software: Pierre-Louis Bazin and Julia Huck; Writing - original draft preparation: Stéfanie A. Tremblay; Visualization: Stéfanie A. Tremblay; Writing - review and editing: Anna-Thekla Jäger, Pierre-Louis Bazin, Chiara Giacosa, Christopher J. Steele, and Claudine J. Gauthier; Funding acquisition: Arno Villringer; Supervision: Christopher J. Steele and Claudine J. Gauthier.

